# Independent component EEG analysis reveals spectral correlates of emotional and cognitive processing in immersive VR environments

**DOI:** 10.1101/2025.11.27.690942

**Authors:** Irene Fondón, María Luz Montesinos

**Author notes:** **Corresponding autor:** María Luz Montesinos, Departamento de Fisiología Médica y Biofísica, Universidad de Sevilla, Av. Sánchez-Pizjuán 4, E-41009 Sevilla, SPAIN.

## Abstract

Virtual reality (VR) is a powerful medium for eliciting emotional states in controlled yet immersive conditions, making it valuable in affective neuroscience and human-computer interaction research. Thus, VR enables multisensory experiences that enhance the sense of presence and immersion of users, which are key factors in modulating affective responses. Building on these capabilities, a recent study developed a protocol for inducing emotions in VR while simultaneously recording electroencephalographic (EEG) activity, with the goal of constructing a database for future research in emotion recognition (Marqués Valderrama et al., 2023). Although the authors provided a detailed description of their experimental design and scenario development, the EEG recordings collected through this protocol remained unanalysed. In the present study, we conduct an in-depth analysis of this dataset. The objective of this study is to characterize the EEG signatures associated with distinct emotional states elicited by immersive VR scenarios and explore their potential for computational modelling in automatic emotion recognition systems.

## 1. Introduction

Understanding the neural dynamics underlying emotional processing remains a central objective in affective neuroscience. Among the available neuroimaging techniques, EEG is particularly well-suited to study emotional states in emotion recognition systems (ER), thanks to its excellent temporal resolution, portability, and sensitivity to rapid cortical fluctuations [1–3]. Moreover, EEG is valued for its inherent objectivity, cost-effectiveness and non-invasive nature. Unlike external manifestations (such as voice or facial expressions), which are susceptible to conscious control, EEG signals reflect authentic neural activity. This provides a more objective and less biased window into an individual’s emotional state.

Emotional experiences have been shown to modulate brain oscillations, with specific frequency bands associated with various aspects of affective and attentional regulation [4,5]. Usually, the characterization of emotional states via EEG has centered on examining variations, at the channel-level, in the spectral power of evoked oscillations within specific frequency bands, especially in the theta (4-8 Hz), alpha (8-13 Hz), and beta (13-30 Hz) ranges, following the presentation of static images. For instance, frontal alpha asymmetries have been associated with emotion valence [6,7], whereas increased beta activity in frontoparietal regions has been associated with high arousal and attentional engagement [4,8]. Thus, arousal is often correlated with increased power in higher frequency bands, especially beta and gamma (>30 Hz), and reduced alpha power [4,9]. Moreover, the beta band power increases in response to low valence [4,8]. In addition, increased theta power is associated with negative valence and high arousal emotions [8]. Additionally, theta activity, especially in midline frontal areas, has been linked to emotional conflict, cognitive load, and salience processing, often increasing in response to emotionally charged stimuli regardless of valence [10,11]. These findings support the fact that emotional engagement can be captured by a distributed pattern of frequency-specific modulations across multiple cortical areas. Moreover, neural oscillations in the alpha and beta ranges can be subdivided into functionally distinct sub-bands. For example, in some studies, alpha activity has been split into low-alpha (8-10 Hz) and high-alpha (10-12 Hz), and the beta band can be divided into beta1 (13-16 Hz), beta2 (16-20 Hz), and beta3 (20-30 Hz) [12,13]. These subdivisions provide a more detailed framework for interpreting frequency-specific power modulations related to affective and cognitive states.

Recent advancements have highlighted VR as an exceptional modality for emotion induction in affective computing research [2,3,14–16]. VR presents a viable trade-off between experimental control and realism [17]. It facilitates the creation of rich, engaging, and contextually detailed computer-generated scenarios [18], enabling the naturalistic and effective elicitation of specific emotional states within a controlled laboratory environment [1,14,17,19]. Several studies confirm that VR environments can induce stronger emotions and a profound sense of immersion and presence compared to traditional two-dimensional (2D) displays [15,20]. This ecological validity provided by immersive VR is vital for obtaining authentic affective responses [20,21].

Despite the promising synergy, integrating EEG acquisition with VR stimuli poses significant technical challenges [15]. The dual use of EEG sensors and VR headsets (HMDs) creates complexity in configuring and synchronizing data flows [1,22,23]. Critically, VR environments inherently introduce noise and artifacts into the EEG signals, since participants are often required to move their heads to fully explore immersive VR environments, which leads to motion artifacts [1,24–26]. These movements, along with blinks, can degrade signal quality, potentially worsening the electrode-scalp connection and causing electrical disturbances [1,15].

Traditional methods for emotion recognition in EEG largely rely on extracting frequency-domain features, such as Power Spectral Density (PSD) and Differential Entropy (DE) [3,19,25], followed by classification using machine learning (ML) or deep learning (DL) algorithms (e.g., SVM, Random Forest, CNN-LSTM hybrid models) [2,15,20,21,27]. While these classification approaches have achieved significant results (often ranging from 73% to 85% accuracy in VR contexts) [15,19,25], they face limitations. Neural Networks (NNs) typically require large datasets and involve high computational costs due to the high dimensionality of EEG data [21]. Furthermore, many ML/DL models operate as “black boxes”, offering high predictive accuracy without providing clear neurophysiological interpretability regarding where and how the emotions arise in the cortex [2]. Analyses focused solely on sensor-level features (e.g., calculating affective indicators based on electrode ratios) often obscure the precise cortical origins of the observed brain activity [21].

To overcome these limitations and gain insightful neurophysiological understanding, the analysis of brain activity must shift from the scalp (sensor-level) to the cortical sources (source-level) [15,21]. Independent Component Analysis (ICA), often implemented via toolboxes like EEGLAB, is the standard technique used extensively for analysing single-trial EEG dynamics and is essential for data preprocessing [15,17,24,25]. ICA excels at isolating and identifying artefactual components (such as those arising from eye movements or muscle activity) and separating them from genuine neural signals, which is critical given the artifact burden in VR recordings [15,20,24,25].

Achieving robust source localization and precise ICA is conventionally predicated on the use of high-density EEG setups [15,16]. The literature emphasizes that higher density electrode configurations (typically 64 channels or more) are required to capture the necessary spatial detail in the brain’s electrical fields, thereby ensuring the reliability of source reconstruction [16]. For instance, dedicated VR emotion datasets, such as the VREED database, utilize 59 high-density EEG channels for this purpose [16,24], and others, like the VRSDEED dataset, employ 128 channels [15]. Conversely, the use of low-density EEG configurations is noted to limit holistic cerebral cortex analysis [15]. Studies employing low-cost, low-channel systems (e.g., 4 channels) often achieve reasonable classification results but typically rely on traditional feature extraction and standard ML/DL classifiers, moving away from detailed source analysis [26].

The current study addresses this gap by focusing on the characterization of emotion using a low-density EEG configuration, specifically, a 14-channel EEG dataset introduced by Marqués Valderrama et al. (2023). This approach directly challenges the prominent assumption in the literature that robust ICA and subsequent source analysis require high-density electrode coverage. The protocol developed by Marques Valderrama et al. established a method for inducing emotions in VR while recording EEG, but the dataset itself remained unanalyzed. Our research, therefore, constitutes the first detailed spectral analysis of these recordings. Using this dataset, we investigate how distinct emotional states, induced in immersive VR, modulate EEG power spectra at the source-level. In contrast to traditional sensor-level analyses, we apply a rigorous source-level methodology, utilizing ICA and IC-clustering to identify consistent neural sources across participants. This work contributes to the field by validating this approach on a low-density setup. By demonstrating that interpretable neurophysiological insights can be reliably obtained from 14-channel data, this research fills a critical gap, paving the way for more accessible and resource-efficient affective computing applications in immersive VR environments.

## 2. Methods

### 2.1. EEG data

The EEG participant data were obtained under a VR EEG Dataset EULA agreement and consisted of pre-processed data (in CSV format) following the application of a zero phase-lag bandpass Butterworth filter (0.5-100 Hz) and a zero phase-lag notch Butterworth filter (50 Hz), as previously described [1]. The corresponding Dreamdeck video file for each participant was also provided to facilitate the alignment between the EEG signals and the VR scenarios. The original study included 23 healthy volunteers. However, following a detailed quality assessment of the raw recordings, 5 datasets were excluded because of severe artifacts or incomplete channel data. Thus, the present analysis includes data from 18 participants (7 females; mean age 21.1 ± 2.3 years; 11 males; mean age 21.7 ± 2.7 years).

### 2.2. Preprocessing

Preprocessing and analysis were performed using version v2025.0.0 of the EEGLAB toolbox [28]. EEG data were imported from CSV files using a custom MATLAB (version 2024a, MathWorks, Natick, MA, USA) script and converted into the EEGLAB dataset format. Channels were renamed and localized according to the standard 10-05 system using the standard_1005.elc file provided by EEGLAB. Event markers corresponding to the experimental conditions (Forest, Alien, City, Museum) were inserted based on participant-specific video timing data. Additionally, the EEG segment corresponding to the introductory portion of the VR experience (featuring a white screen and a simple welcome message at the very beginning of the video) was used as a neutral control condition due to its minimal visual and emotional content. Data were bandpass filtered between 1 and 45 Hz to optimize ICA performance. After removing bad channels (one channel in 7 out of 18 datasets) and manually rejecting noisy segments, the data were re-referenced to the average of all remaining channels.

### 2.3. EEG cortical source localization and clustering

ICA was performed using the *runica* algorithm implemented in EEGLAB, to decompose the EEG data into independent components characterized by fixed spatial patterns. These components included sources related to neural activity and artifacts, such as blinks, eye movements, and muscle activity. Following ICA, equivalent current dipoles were estimated for each component using the DIPFIT plugin [29], based on a standardized three-shell boundary element model derived from the Montreal Neurological Institute (MNI) template brain and included in EEGLAB. Dipole fitting was performed using the EEGLAB automated settings. For subsequent group-level analyses, the data were segmented by condition (Neutral, Forest, Alien, City, Museum) for each participant. Only components whose estimated dipoles were located within brain volume and whose residual variance (RV) was below 15% (indicating that the dipole explained at least 85% of the scalp projection) were retained for further analyses [30].

For group-level IC-clustering, preclustering was conducted using the EEGLAB STUDY framework. The features used for preclustering included equivalent dipole location (PCA-reduced to 3 dimensions; weight = 1), dipole orientation (PCA-reduced to 3 dimensions; weight = 0.3), and scalp topography (PCA-reduced to 13 dimensions, corresponding to the number of EEG channels minus one; weight = 1). Clustering was performed using the Optimal k-means algorithm, exploring a range of 5-15 clusters to automatically determine the optimal solution. Outlier ICs were excluded based on a criterion of exceeding 3 standard deviations from any cluster centroid.

### 2.4. Spectral power analysis

For each IC cluster, the spectral power was extracted using the fast Fourier transform (FFT) implemented within the EEGLAB STUDY framework. The spectral analysis (4-40 Hz) focused on the theta, alpha, beta, and low-gamma frequency bands. The delta band (1-4 Hz) was excluded because of its high susceptibility to low-frequency artifacts. Frequencies above 40 Hz were also excluded to reduce contamination from high-frequency muscle artifacts, which could be particularly prevalent in VR EEG recordings. This restriction necessarily precluded the analysis of the high gamma band (>50 Hz), a frequency range increasingly implicated in localized emotional and cognitive processing in VR environments [31].

The spectral regions showing significant differences between conditions (p < 0.05) were identified using the paired permutation statistics method in EEGLAB (200,000 permutations), with false discovery rate (FDR) correction applied to control for multiple comparisons.

## 3. Results

### 3.1. Description of the original experimental protocol

In the original experimental setup described by the authors [1], VR environments were employed to elicit distinct emotional responses within a relatively short time frame while minimizing user interaction to help reduce motion-related artifacts in the EEG recordings.

The experimental protocol involved two VR experiences. The first one was a guided tutorial using the Google Earth VR application (with a duration of approximately 5 min) to familiarize the participants with VR. The second experience was the Oculus Dreamdeck VR experience (with a duration of approximately 3 min), comprising a sequence of four brief, immersive scenarios selected by the authors to elicit distinct emotional responses. These were: Lowpoly Forest (34 s), designed to evoke relaxation via a stylized natural landscape with soft ambient sounds and minimal sensory stimulation; Alien (33 s), intended to induce excitement by placing the participant near a humanoid alien figure speaking in an unintelligible fictional language within an unfamiliar, extraterrestrial environment; Futuristic City (24 s), aimed to elicit surprise through an aerial view of an urban skyline; and T-Rex (49 s), intended to provoke fear by simulating the sudden approach of a dinosaur within a museum obscure corridor. Notably, none of the Dreamdeck scenarios involved the interaction of the user with controllers, reinforcing a passive observational role.

A traditional theoretical framework used to describe emotion in cognitive neuroscience is the Circumplex Model of Affect [32], which maps emotional states onto a two-dimensional space defined by valence (pleasant-unpleasant) and arousal (high-low). This continuous representation enables the description of small affective changes without using strict category labels. In the original study by Marqués Valderrama et al. (2023), the emotional responses were evaluated through a Self-Assessment Manikins (SAM) approach, rating both valence and arousal for each scenario from 1-7. The Forest condition returned a relatively high valence (mean = 5.75; SD = 0.87) and low arousal (mean = 1.83; SD = 1.47), indicative of a relaxed and positive emotional state. The Alien scenario resulted in moderate valence (mean = 4.33; SD = 1.72) and high arousal (mean = 4.33; SD = 1.23), reflecting excitement or alertness. The City scenario provoked a moderately high valence (mean = 5.08; SD = 1.68) and a relatively high arousal (mean = 4.25; SD = 0.87), suggesting surprise or engagement. Finally, the Museum (T-Rex) scenario showed the lowest mean valence (mean = 4.25; SD = 2.49) and the highest arousal (mean = 5.75; SD = 0.87), consistent with fear- or threat-related emotional state.

### 3.2. IC clusters

The original EEG recordings from the Dreamdeck experience were provided in CSV format and were processed as described in the Methods section. In brief, the preprocessing pipeline included bandpass filtering, manual artifact rejection, ICA decomposition, equivalent dipole fitting, and IC-clustering. As previously described, the duration of the raw EEG data varied across conditions, ranging from 24 to 49 s. Following manual cleaning, the mean retained duration per condition ranged from 17.1 to 39.6 s, reflecting a data loss of approximately 19%-29% depending on the condition (see Supplementary Table S1).

A total of 227 ICs were initially extracted across the 18 participants included in the study. This figure was reduced to 116 ICs when only the dipoles located within the brain and showing RV below 15% were selected. The clustering algorithm identified 5 clusters and 2 outlier ICs. An additional outlier IC, showing an atypical scalp topography, was manually removed. The cluster B (see below) included 19 ICs clearly classified by the IClabel EEGLAB plugin as ocular artifacts (“eye”), whereas the cluster C contained one “eye” IC. Eye-related and outlier ICs were manually removed before further analyses. After this cleaning, 93 ICs remained, corresponding to approximately 5 ICs per subject across the 18 participants, which is consistent with values reported in the literature for EEG studies using ICA and dipole fitting.

The IC clusters obtained (Table 1 and Figure 1 and 2) were localized in the left secondary visual cortical region (cluster A; 20 ICs from 15 participants), left anterior prefrontal region (antPFC) (cluster B; 18 ICs from 12 participants), right dorsolateral prefrontal region (DLPFC) (cluster C; 20 ICs from 15 participants), left angular gyrus region (cluster D; 14 ICs from 12 participants), and right fusiform gyrus region (cluster E; 21 ICs from 13 participants).

**Figure 1.**
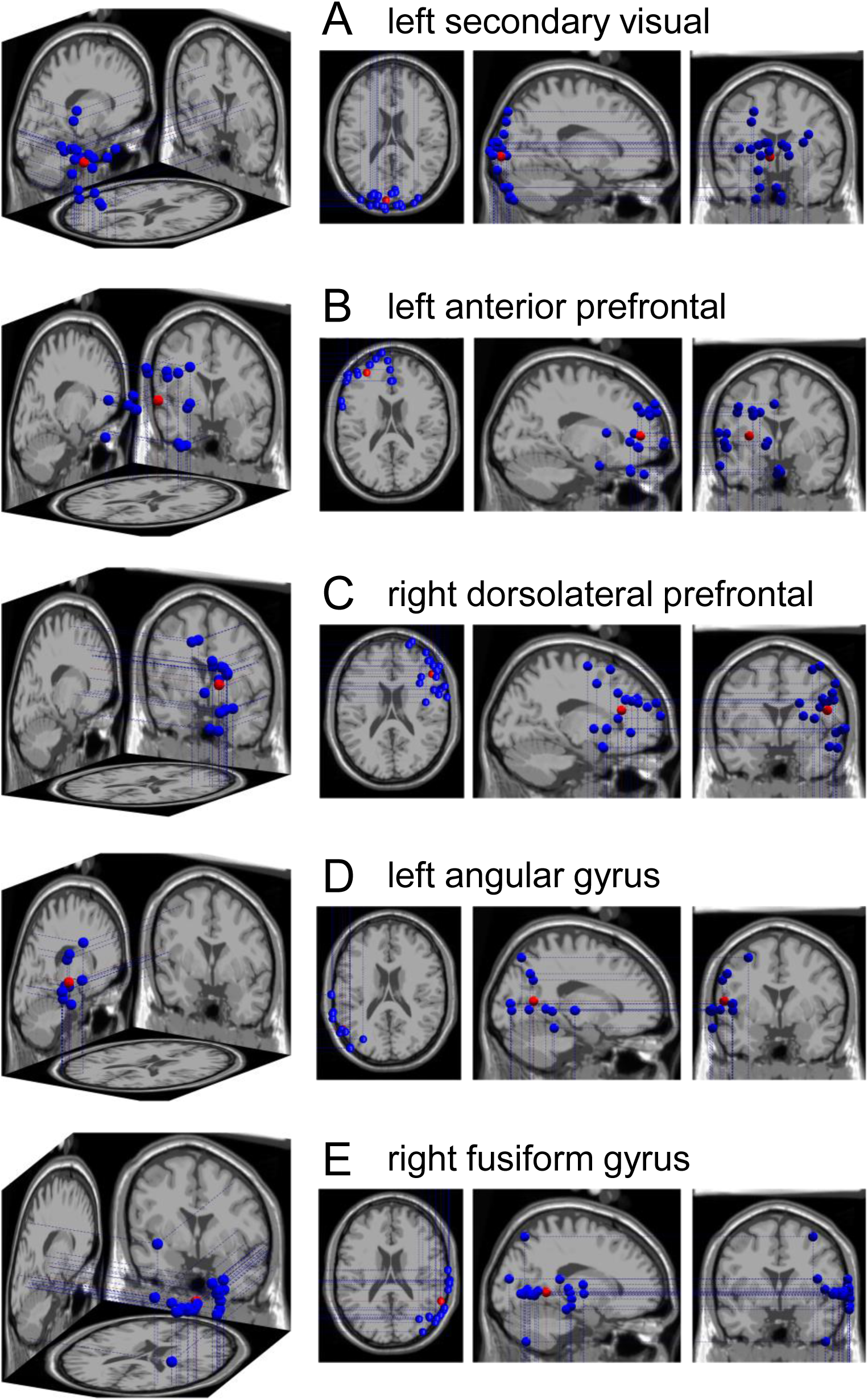
Dipole localization of IC clusters. Clusters A-E are visualized in the Montreal Neurological Institute brain volume in top, sagittal, and coronal views. The blue points indicate the dipole locations for each IC included in this study. and the red points indicate the corresponding cluster centroid.

**Figure 2.**
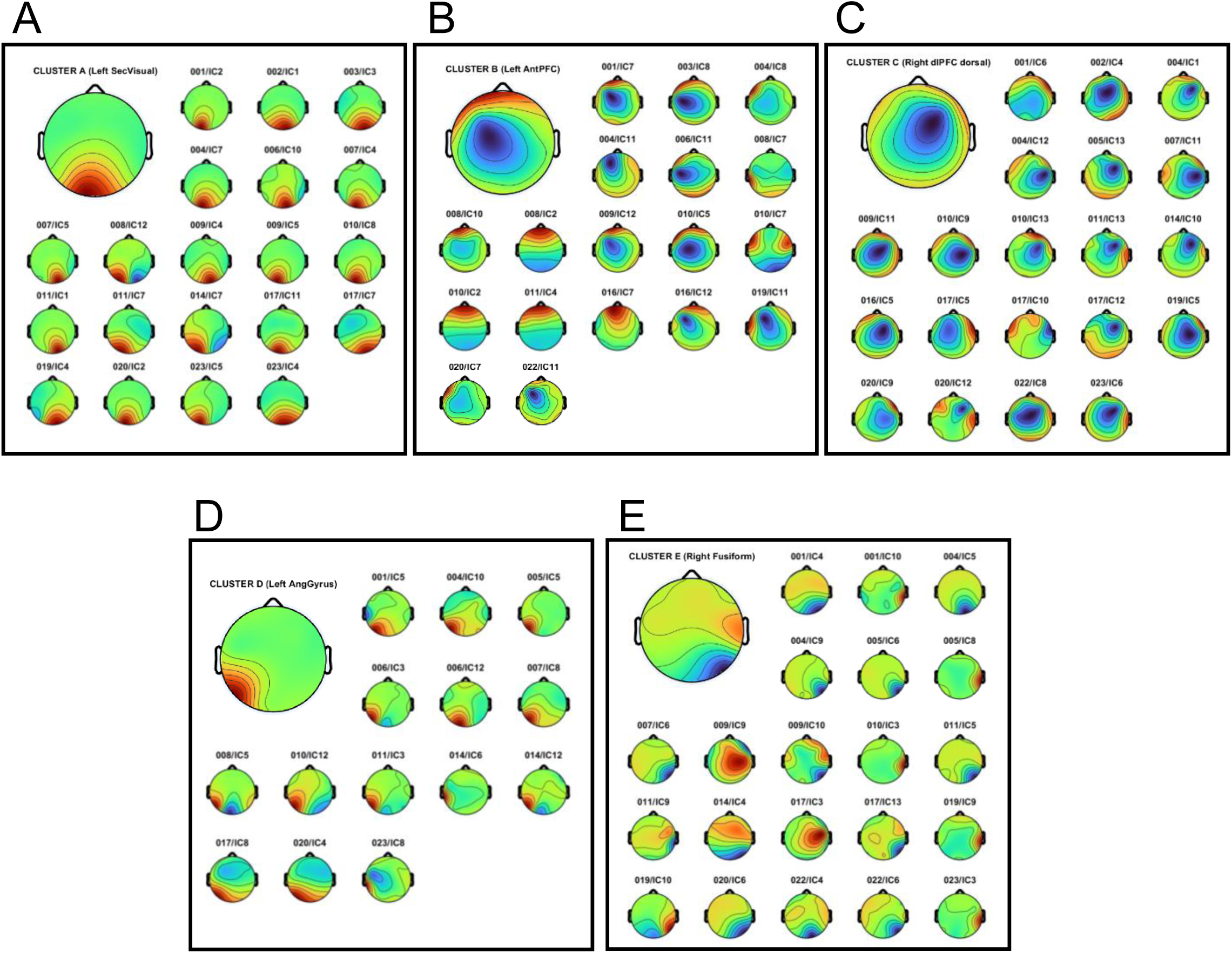
Scalp maps of IC clusters. The individual and the mean scalp maps averaged across components included in each cluster are shown. The original subject identifier (001-023) and the corresponding IC numbers are shown for each individual scalp map.

**Table 1.** Clusters of independent sources obtained by ICA.

| Cluster | Location of the cluster<br>centroid | Cluster centroid<br>coordinates<br>(x, y, z) | Number of<br>participants<br>(Ps) and ICs |
| --- | --- | --- | --- |
| A | left secondary visual cortical<br>region (BA18) | -7.2, -95.9, -2.4 | 15 Ps, 20 ICs |
| B | left anterior prefrontal region<br>(BA10) | -31.2, 47.9, 6.2 | 12 Ps, 18 ICs |
| C | right dorsolateral prefrontal<br>region (BA9) | 50.1, 29.5, 19.1 | 15 Ps, 20 ICs |
| D | left angular gyrus cluster<br>(BA39) | -57.3, -61.8, 9.7 | 12 Ps, 14 ICs |
| E | right fusiform gyrus cluster<br>(BA37) | 62.3, -48.7, 0.7 | 13 Ps, 21 ICs |

### 3.3. Spectral analysis of the IC clusters

The spectral power for each cluster and condition was evaluated from 4 to 40 Hz, and the statistically significant differences across the four VR scenarios (Forest, Alien, City, Museum) and the Neutral control condition were determined.

In the left secondary visual region (cluster A; Figure 3), which is associated with visual processing and perceptual integration, the Forest condition elicited significantly greater power than the Neutral condition in the beta3 band (23-28 Hz) and low-gamma (34-35, 39-40 Hz) bands, suggesting enhanced visual engagement. The City condition showed higher power than the Neutral condition in theta (4-9 Hz) and throughout the beta and low-gamma range (13-40 Hz). Compared with Forest, City exhibited greater power in beta2 (15-20 Hz), beta3 (23-27, 29-30 Hz), and low-gamma (31-32, 34-36, 38 Hz) bands. Moreover, City elicited significantly greater power than both Alien and Museum across the entire 4-40 Hz spectrum.

**Figure 3.**
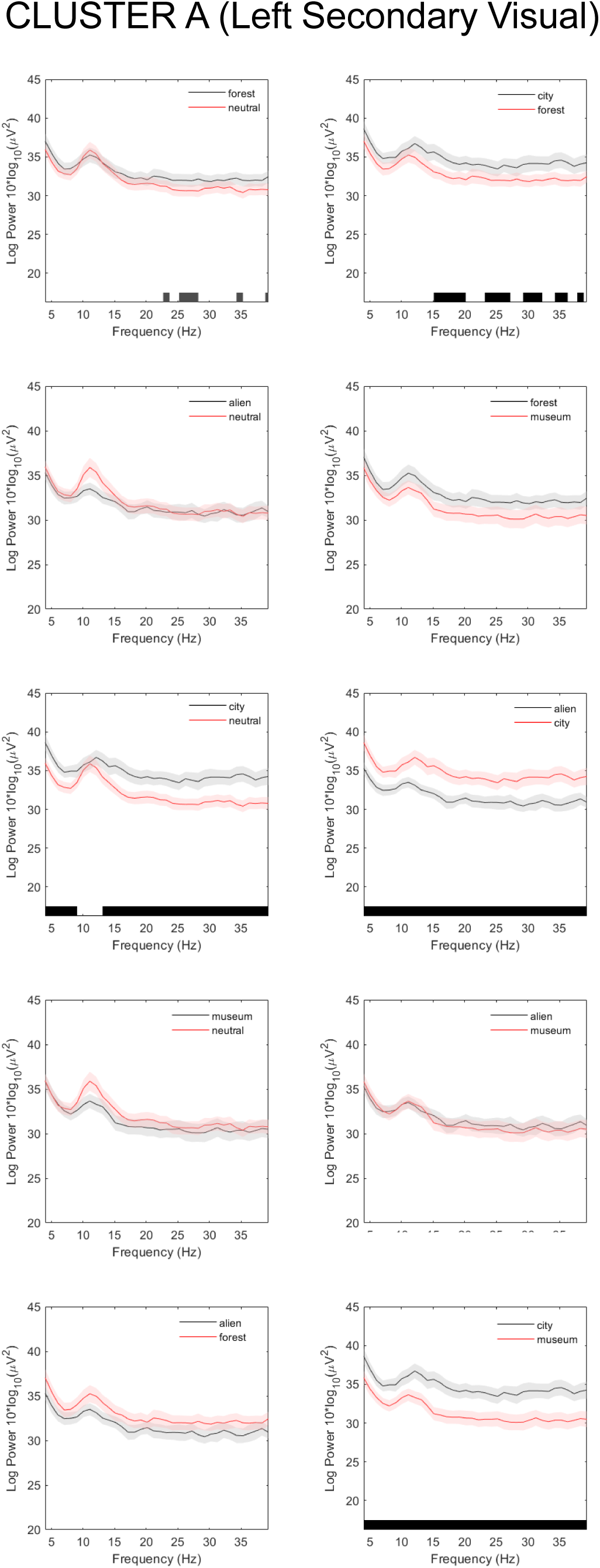
Power spectrum of IC cluster A (left secondary visual region). The average power of all ICs included in the cluster (lines) and the corresponding standard error (shading) are shown for each pair of conditions. The black underbars denote spectral regions of statistically significant differences (FDR-corrected p value < 0.05).

In the left antPFC region (cluster B; Figure 4), a region implicated in emotional regulation and decision-making [33,34], Museum elicited higher power than Neutral from 14-40 Hz, encompassing the beta and low-gamma bands. Compared with Forest, Museum showed greater power in beta2 (17-20 Hz), beta3 (24-26 Hz), and low-gamma (28-40 Hz) bands, whereas Forest exceeded Museum at 5 Hz (theta). Museum also exhibited higher power than Alien at 31, 35, and 39 Hz, and surpassed City in beta2 (18-20 Hz), beta3 (24-26, 29 Hz), and low-gamma (33-38 Hz) bands, reflecting increased high-frequency activity.

**Figure 4.**
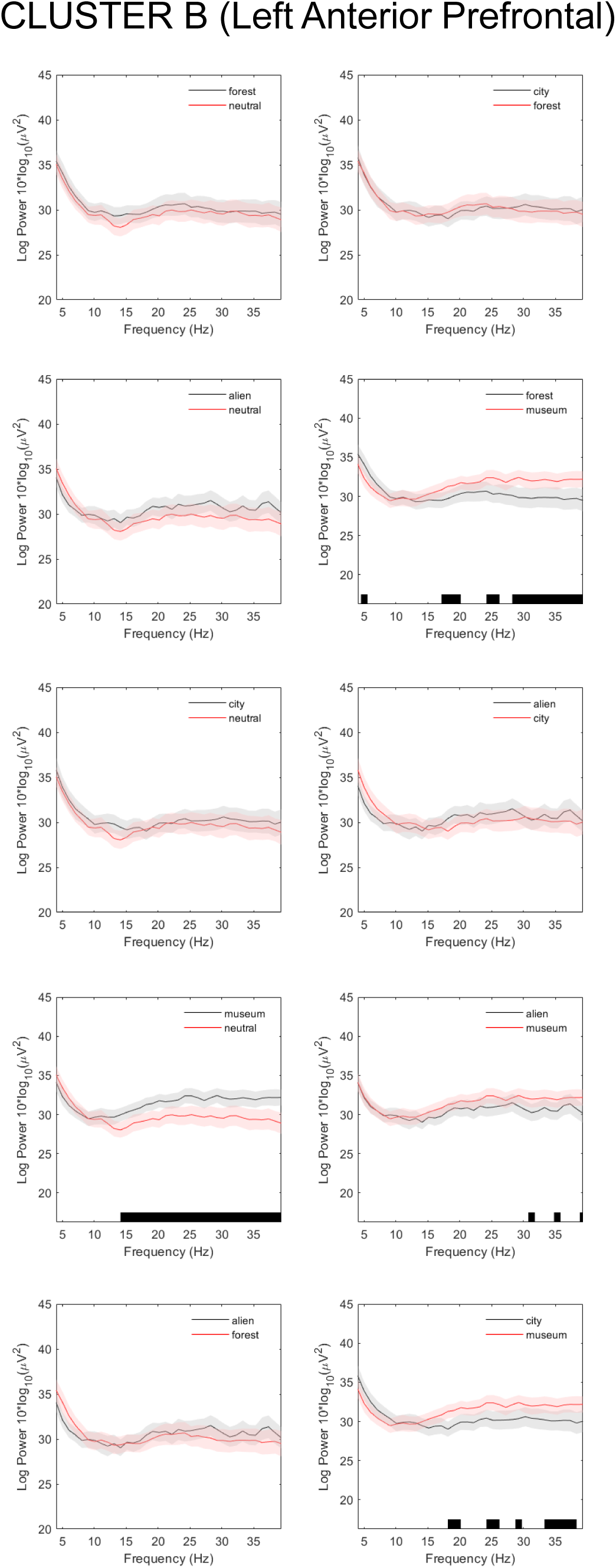
Power spectrum of IC cluster B (left anterior prefrontal region). The average power of all ICs included in the cluster (lines) and the corresponding standard error (shading) are shown for each pair of conditions. The black underbars denote spectral regions of statistically significant differences (FDR-corrected p value < 0.05).

In the right DLPFC region (cluster C; Figure 5), which is involved in executive functions, decision-making, and top-down emotional modulation [34–37], the only statistically significant difference was observed between the Museum and Forest scenarios. The Museum condition elicited greater spectral power than the Forest condition across a wide frequency range (12-40 Hz), encompassing the upper alpha, beta, and low-gamma bands.

**Figure 5.**
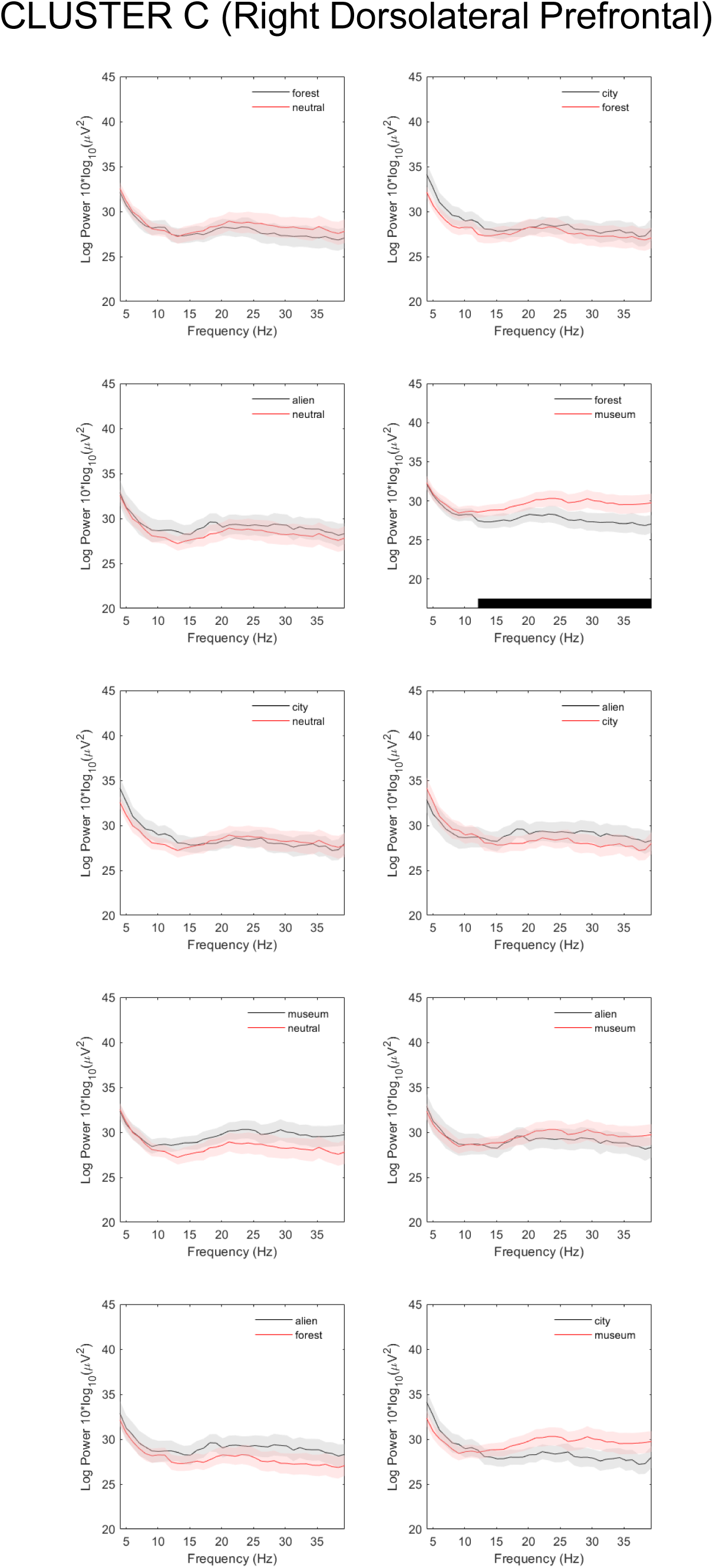
Power spectrum of IC cluster C (right dorsolateral prefrontal region). The average power of all ICs included in the cluster (lines) and the corresponding standard error (shading) are shown for each pair of conditions. The black underbars denote spectral regions of statistically significant differences (FDR-corrected p value < 0.05).

In the left angular gyrus cortical cluster (cluster D; Figure 6), associated with semantic integration, attention, and visuospatial processing [38,39], Forest exhibited greater power than Alien at 11 Hz (alpha). The City condition provoked a significantly higher power than the Neutral condition across the full 4-40 Hz spectrum. Compared with Forest, City showed greater power at 4 Hz and 9-40 Hz. It also exceeded Alien in theta (4, 6 Hz), alpha (8-13 Hz), beta2-3 (15-30 Hz), and low-gamma (31-37, 39 Hz) bands. Finally, compared to Museum, City showed higher power at 4 Hz, 6-7 Hz (theta), and 9-10, 12 Hz (alpha). These findings suggest robust and widespread activation of the left angular gyrus cluster in the City scenario.

**Figure 6.**
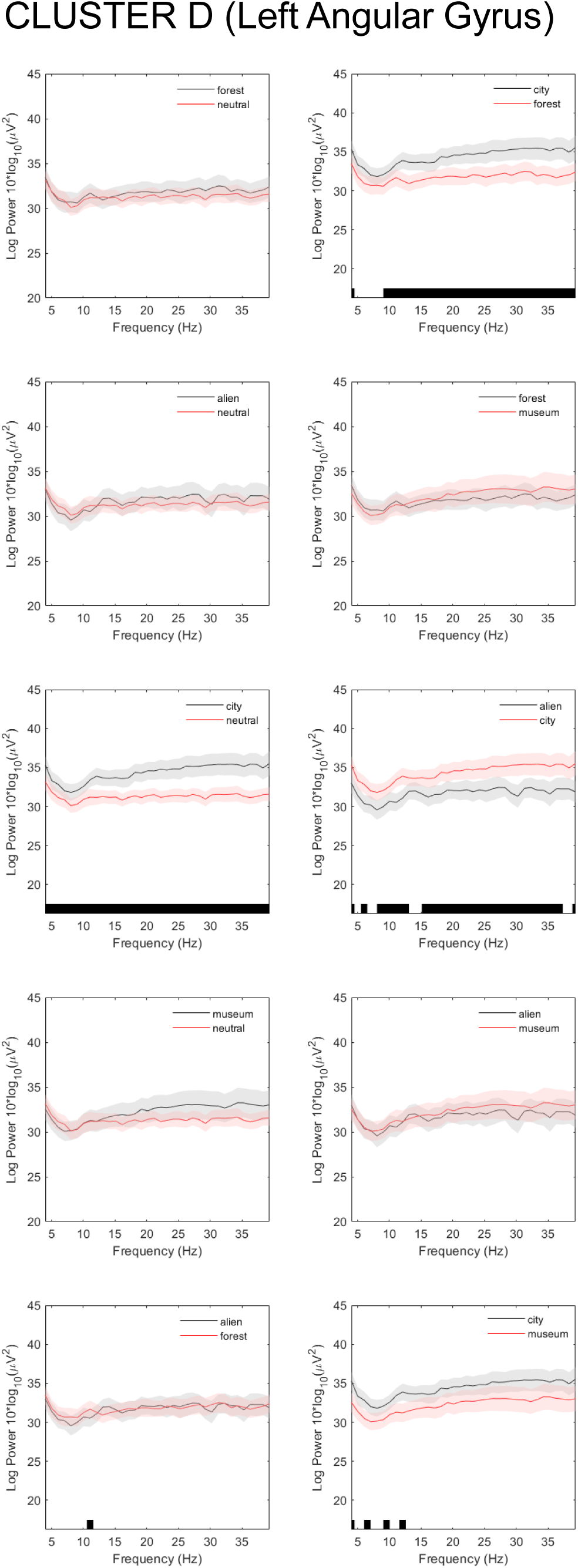
Power spectrum of IC cluster D (left angular gyrus cluster). The average power of all ICs included in the cluster (lines) and the corresponding standard error (shading) are shown for each pair of conditions. The black underbars denote spectral regions of statistically significant differences (FDR-corrected p value < 0.05).

Finally, in the right fusiform gyrus region (cluster E; Figure 7), known for its role in visual object recognition and affective face and language processing [40–43], spectral modulations were observed in the lower frequency bands. The Forest scenario elicited greater power than the Alien condition at 6 Hz (theta), whereas the City condition showed significantly higher power than the Alien scenario from 4 to 6 Hz and exceeded the Museum condition at 4 Hz.

**Figure 7.**
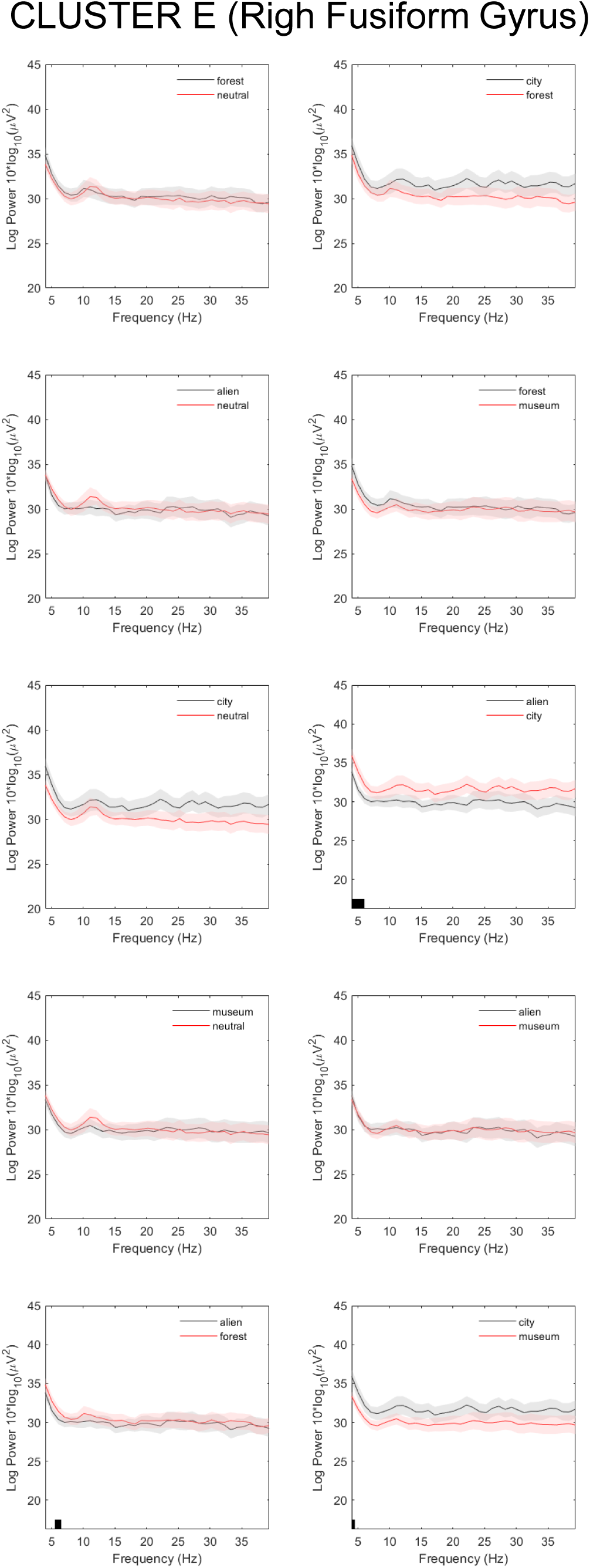
Power spectrum of IC cluster E (right fusiform gyrus cluster). The average power of all ICs included in the cluster (lines) and the corresponding standard error (shading) are shown for each pair of conditions. The black underbars denote spectral regions of statistically significant differences (FDR-corrected p value < 0.05).

## 4. Discussion

In this study, we investigated the spectral changes of EEG IC clusters during a continuous VR experience aimed at eliciting different emotions, revealing condition-specific modulations in distributed cortical regions associated with visual, cognitive, and affective processing. Although source-based EEG analyses usually rely on high-density EEG systems (more than 32 channels), previous work has shown that meaningful separation of neural components and cross-subject clustering can be achieved even with reduced channel counts [44], suggesting that low-density EEG systems, as is the case here, can still support advanced signal decomposition and source-level functional interpretation when appropriate ICA preprocessing is applied.

### 4.1. Visual engagement and the influence of stimulus complexity and temporal order

The spectral pattern obtained for the left secondary visual region (Cluster A) in the different conditions points to City as the scenario that elicited the strongest neural engagement in visual regions, likely reflecting its immersive, dynamic, and visually rich content. Although Forest also showed increased high-frequency (beta3 and low-gamma) power compared to Neutral, the absence of differences between Forest and Alien, despite the latter being more visually complex, suggests that stimulus order may have shaped the neural responses. Given the fixed sequence of conditions (Neutral, followed by Forest, then Alien, then City, and, finally, Museum), it is plausible that the initial increases in visual attention and arousal during Forest led to partial habituation, dampening the neural response to subsequent scenarios like Alien and even Museum. The lack of significant differences between Museum and Neutral or Museum and Forest, for example, despite its perceptual complexity, supports this interpretation. These findings suggest that both perceptual complexity and temporal context influence cortical engagement during immersive VR experiences.

### 4.2. High-frequency prefrontal region engagement in the Museum condition

The increased beta and/or low-gamma power observed in the left antPFC region (Cluster B) during the Museum condition, compared to all the other scenarios, likely reflects high engagement of cognitive and/or emotional regulation processes in response to the intense and potentially threatening content of this scenario [4]. Specifically, increased beta and low-gamma activity in this area is a robust marker of executive functions, top-down control, and elevated cognitive load or demand [45], which would be expected during threat assessment (i.e., Museum condition). The antPFC is known to be involved in internally focused thoughts. It plays a key role in monitoring and evaluating the outcomes of earlier cognitive processes, helping individuals to assess their own thoughts, feelings, and decisions [33,34]. Thus, the Museum condition showed greater beta2 and beta3 activity than Neutral, Forest, and City, consistent with the proposed role of beta oscillations in top-down control and affective engagement [46–48]. Interestingly, the differences were observed only in the low-gamma range when comparing Museum to Alien. This suggests that, while both conditions similarly evoked a rather negative emotional valence, the stronger arousal associated with Museum may have driven greater recruitment of regulatory and integrative mechanisms [49]. Thus, this enhanced gamma power could be a neural correlate of threat evaluation and anticipatory response management under higher emotional intensity [50].

The Museum condition also exhibited higher spectral power than the Forest condition in the right DLPFC region (Cluster C) across a broad frequency range (12-40 Hz), encompassing high-alpha, beta, and low-gamma bands. This finding also supports the relevance of the prefrontal region for managing the complex content of the Museum scenario, probably engaging mechanisms involved in immediate executive control and top-down modulation [34,37,51].

### 4.3. Multimodal integration and attentional demands in the left angular gyrus cluster in the City scenario

During the City scenario, the left angular gyrus cluster showed widespread increases in spectral power across theta, alpha, beta2-3, and low-gamma bands relative to other conditions. This pattern aligns with the established role of the angular gyrus in multimodal integration, attention, and visuospatial cognition [39,52]. Although alpha and theta oscillations are often linked to relaxed states, enhanced power in these bands within task-relevant cortical areas, such as the angular gyrus, has been interpreted as reflecting active top-down attentional control and functional inhibition, facilitating the processing of complex stimuli [53,54]. Accordingly, the City scenario, characterized by a visually rich, futuristic landscape devoid of linguistic or social content, likely demanded a sustained attentional engagement and integrative processing of novel spatial features. The increased beta and low-gamma power observed in City compared to Neutral, Forest, and Alien further suggests increased cognitive and sensory integration efforts, in consonance with the association of beta and gamma bands with active information processing and attention maintenance [49,50]. In contrast, despite its visual complexity and emotional intensity, the absence of similar beta and low-gamma increases in the Museum condition may indicate differential processing demands: the Museum scenario likely engages more executive and regulatory circuits in prefrontal areas for emotional control, with less reliance on angular gyrus-mediated visuospatial and multisensory integration. Additionally, the fixed order of scenarios and potential habituation effects (already discussed) might have attenuated integration demands in later conditions, such as Museum.

The elevated alpha power in the left angular gyrus cluster (also involved in semantic processing) during the Forest condition compared to the Alien scenario may initially appear paradoxical, considering the lower visual complexity of Forest and its naturalistic animal sounds versus the more complex visual and auditory stimuli characteristic of the Alien environment, which includes an invented language. However, as discussed before, alpha oscillations are widely recognized not only as indicators of relaxation but also as a reflection of active top-down inhibitory control mechanisms that suppress task-irrelevant information to facilitate selective attention [53]. Conversely, the Alien scenario, with its unfamiliar multisensory content, likely elicits alpha desynchronization (i.e., power reduction) due to increased cortical excitation and integrative demands [55].

### 4.4. Modulation of the theta band across conditions in the right fusiform gyrus

As previously mentioned, the right fusiform gyrus is involved in visual object recognition and affective face and language processing [41–43,56]. This region has also been associated with functional connectivity to the angular gyrus for integrating shape and colour information. Interestingly, the Forest scenario elicited higher theta power than Alien, although Alien presented more complex social stimuli, such as faces and an invented language. In fact, the Alien scenario did not elicit significantly greater spectral power in any frequency band across the cortical regions examined. This absence of localized enhancement, despite the moderate valence and high arousal, as well as high multisensory complexity, of the Alien condition may reflect a more diffuse or less coherent neural response to its ambiguous and unfamiliar features. The fixed temporal position of Alien (following Forest) may have compounded these effects, with the sudden increase in novelty potentially causing transient disorientation.

The City scenario elicited greater theta power than both Alien and Museum in this cluster, that could be interpreted as enhanced perceptual integration and attentional demands in processing novel, non-social stimuli [57]. This pattern might reflect the sensitivity of the fusiform gyrus not only to socially and affectively relevant visual cues, such as faces, but also to complex object features that may require detailed perceptual analysis.

### 4.5. Summary of the spectral changes considering the temporal dimension of the different scenarios

The sequential analysis of the spectral dynamics across the scenarios revealed a progression of functionally distinct cortical modulations (Figure 8). Thus, the only significant change from the Neutral scenario to the Forest condition was an increase in beta3 and low-gamma power in the left secondary visual region, likely reflecting enhanced visual engagement with the Forest content. The subsequent transition from Forest to Alien showed selective decreases in alpha power (11 Hz) in the left angular gyrus region and theta power (6 Hz) in the right fusiform gyrus cluster, suggesting a shift from calm, top-down attentional states to increased sensory and semantic disorganization in response to the Alien scenario’s unfamiliar and ambiguous content. Notably, the change from Alien to City marked a broad recruitment of integrative and perceptual areas, with increased power across theta, alpha, beta2-3, and low-gamma bands in the left angular gyrus cluster, additional theta increases in the right fusiform gyrus region, and a generalized 4-40 Hz power increase in the left secondary visual region, indicating intensified attentional, multisensory, and spatial processing driven by the immersive City environment. Finally, the transition from City to Museum was characterized by a redistribution of spectral activity: a decrease of power across 4-40 Hz in the left secondary visual region, increased beta2-3 and low-gamma power in the left antPFC region, and concurrent decreases in theta and alpha bands in the left angular gyrus cluster and theta in the right fusiform gyrus cluster. These changes suggest a shift away from visuo-perceptual integration toward higher-order regulatory and evaluative mechanisms in response to the emotionally charged content of the Museum scenario. These sequential shifts emphasize the importance of considering temporal ordering and scenario transitions when interpreting neural responses during continuous, ecologically valid affective experiences.

**Figure 8.**
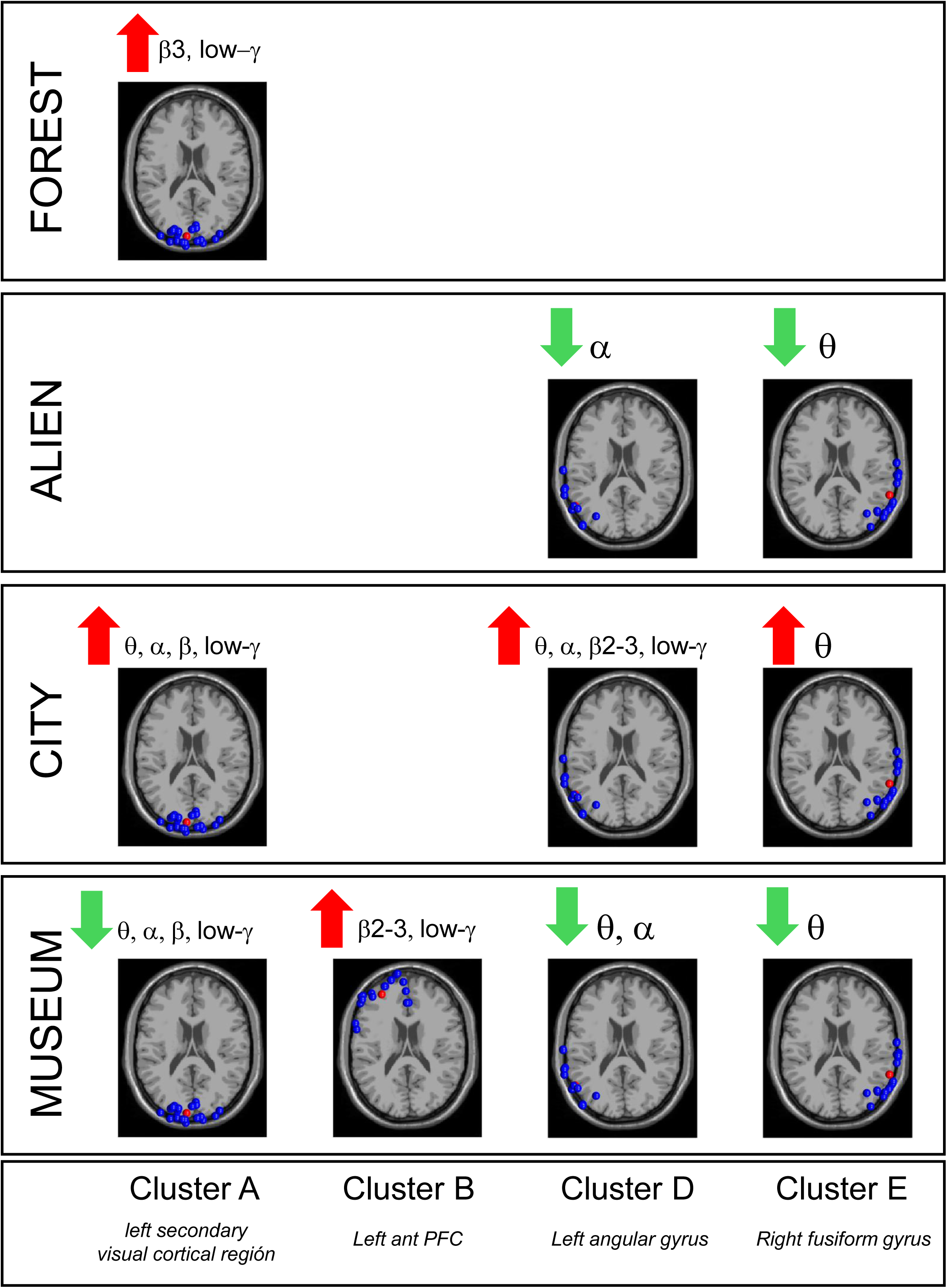
Summary of the spectral changes considering the temporal dimension of the different scenarios. The power spectrum changes for the different clusters, in the Forest scenario, compared to Neutral, in the Alien scenario, compared to Forest, in the City scenario, compared to Alien, and in the Museum scenario, compared to City, are summarized.

### 4.6. Effects on specific beta sub-bands

Across clusters, significant spectral differences were concentrated in the upper beta range, particularly in beta2 (16-20 Hz) and beta3 (20-30 Hz). These effects were consistently observed in the visual (Cluster A), frontopolar (Cluster B), and angular (Cluster D) regions and were associated with emotionally and cognitively demanding scenarios, notably City and Museum. In contrast, beta1 (13-16 Hz), although occasionally included in broader power increases, did not show distinct or selective modulations, suggesting a more limited role in the emotional and attentional processes captured in this study. This frequency-specific profile aligns with prior evidence showing that beta2 and beta3, but not beta 1 power increase with higher cognitive load and executive demands and in response to emotionally salient stimuli [12,47,48].

### 4.7. The role of VR scenario design and immersion in eliciting affective correlates

VR is not merely a stimulus delivery tool, but it constitutes a context generator, playing a central role in eliciting emotion. Affective neuroscience has historically relied on static 2D stimuli (such as the IAPS database) which often lack the ecological validity required to elicit authentic emotional responses. Our findings, which reveal clear and coherent cortical patterns (such as those observed in the prefrontal cortex and angular gyrus), strongly suggest that the immersive VR scenarios were successful precisely because they induced genuine affective states. The ability of VR to immerse the participant, generating a sense of presence, is likely a key amplifying factor that cannot be understated.

From a VR design perspective, the different scenarios (Forest, Alien, City, Museum) varied not only in their intended emotional content (valence/arousal) but, critically, in their cognitive and sensory load. Immersion is not a monolithic concept; it relies on perceptual fidelity and interactivity. It is plausible that the success of our 14-channel analysis, a surprising engineering finding, is partly due to the high-fidelity VR generating stronger, less ambiguous neural responses than would be elicited by simpler stimuli. VR, therefore, does not just present the stimulus; it primes the brain for a more robust response. Nevertheless, from the standpoint of VR research and development, it is essential to recognize an intrinsic limitation of the technology: the potential emergence of cybersickness and other VR-specific perceptual artifacts. While VR induces strong emotions, it can also induce nausea, visual fatigue, or disorientation, all of which contaminate the EEG signal. The very weight of the HMD introduces muscular artifacts (EMG) from the neck and jaw, which are a challenge for any component analysis (especially with only 14 channels). Future studies must include objective and subjective measures of cybersickness to correlate with EEG data and discern whether some cortical activations are related to physical discomfort rather than the target emotion.

### 4.8. Methodological considerations and study limitations

Although the findings of this study are robust, several methodological considerations are worth noting. As mentioned, the VR experience was continuous and the scenarios were presented in a fixed order, which introduces significant possible confounds related to habituation, fatigue, or carry-over effects across conditions. The absence of counterbalancing in the scenario order limits the ability to fully dissociate the sequence effects from true condition-specific differences. Future studies should consider the randomization of the scenario order or the separation of conditions in different blocks.

The unequal durations of the scenarios and the variable lengths of usable EEG data following manual artifact rejection are another limitation of this study. Although this could theoretically bias power estimates obtained through FFT, the existence of significant condition effects in some clusters but not others, most notably the strong responses elicited by City, despite its shorter duration and greater data loss, suggests that such biases did not systematically distort the results. Nevertheless, harmonizing scenario durations and improving EEG quality would strengthen future analyses.

Finally, while the use of ICA-based clustering and source localization supports the functional interpretation of results, the use of a standard MNI head model for the participants and, as already discussed, a relatively low-density EEG montage limits the precision of the anatomical localization of the dipoles. These factors may introduce some localization error [30]. Therefore, anatomical labels should be interpreted with caution and understood as approximate. The cost of using 14 channels is, unequivocally, the spatial precision.

Looking forward, these limitations define clear research directions: (1) Hardware: while for precision research more is better, our study suggests that for accessible commercial or wearable applications, the industry should focus less on adding channels and more on optimizing the position of those 14-20 channels (perhaps based on the stable cluster locations we found). (2) Generalization: while this principle should generalize to other low-density systems, we caution that dry electrodes introduce a different challenge. Their high and variable impedance noise might not be handled as effectively by a basic ICA pipeline, requiring more advanced preprocessing algorithms. (3) Software (machine learning and sources): the obvious next step is to use the cortical sources (ICs) as input features for ML. Instead of feeding ML with “dirty” sensor data, we propose feeding it with the clean output of the clusters (e.g., “power in the frontal cluster”). This would finally create neurophysiologically informed ML models that are robust, interpretable, and viable even on low-density wearable systems.

## CRediT authorship contribution statement

**Irene Fondón:** Conceptualization, Resources, Writing – original draft, Writing – review & editing

**María Luz Montesinos:** Data curation, Formal analysis, Visualization, Writing – original draft, Writing – review & editing

## Declaration of generative AI and AI-assisted technologies in the manuscript preparation process

To enhance the linguistic and stylistic quality of the manuscript, the institutional versions of Trinka AI, NotebookLM and Google Gemini were used, available through agreements between these platforms and the University of Seville. After using these tools, the authors reviewed and edited the content as needed and take full responsibility for the content of the published article.

## Supplemental Data

**Table S1.**
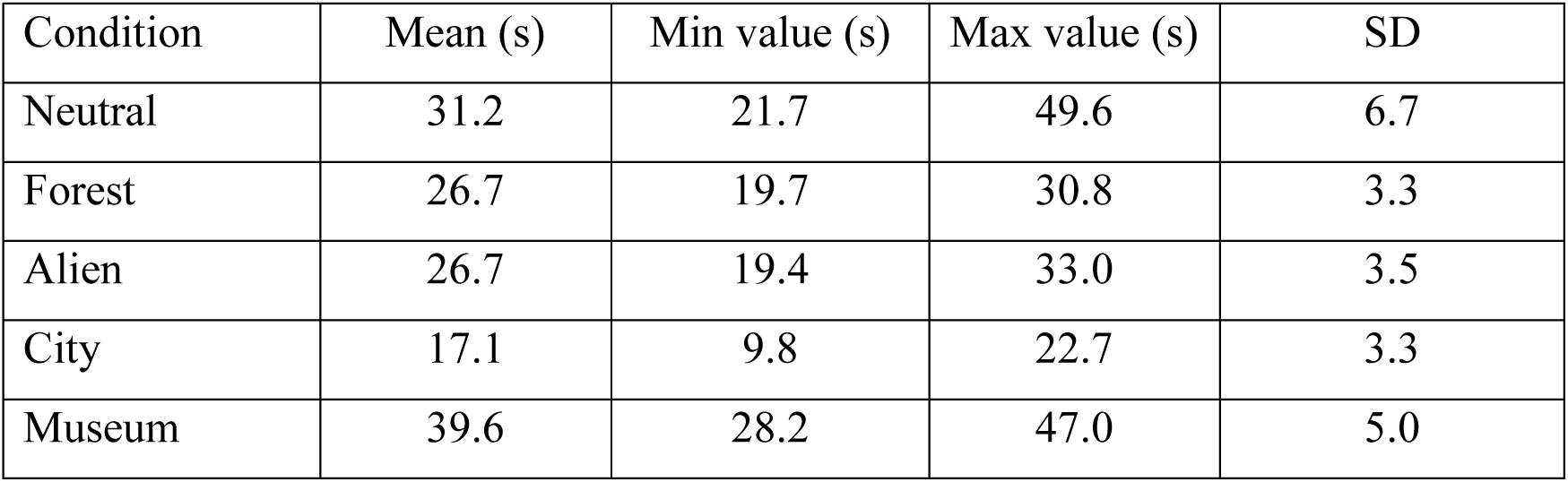
Mean, range, and standard deviation of dataset durations per condition post-cleaning.

